# DNA metabarcoding reveals adaptive seasonal variation of individual trophic traits in a critically endangered fish

**DOI:** 10.1101/2021.01.25.428043

**Authors:** Kurt Villsen, Emmanuel Corse, Emese Meglécz, Gaït Archambaud-Suard, Hélène Vignes, Alexander V. Ereskovsky, Rémi Chappaz, Vincent Dubut

**Author notes:** Correspondence: Kurt Villsen,; or Vincent Dubut.

## Abstract

Dietary studies are critical for understanding foraging strategies and have important applications in conservation and habitat management. We applied a robust metabarcoding protocol to characterize the diet of the critically endangered freshwater fish *Zingel asper* and conducted modelling and simulation analyses to characterize and identify some of the drivers of individual trophic trait variation in this species. We found that intra-specific competition and ontogeny had minor effects on the trophic niche of *Z. asper*. Instead, our results suggest that the majority of trophic niche variation was driven by seasonal variation in ecological opportunity (in our case, the seasonal variation in the availability of preferred prey types). Overall, our results are in line with the optimal foraging theory and suggest that *Z. asper* is specialized on a few ephemeropteran prey species (*Baetis fuscatus* and *Ecdyonurus*) but adapts its foraging by becoming more opportunistic as its favoured prey seasonally decline. Despite the now widespread usage of metabarcoding, very few studies have attempted to study inter- and intra-populational individual trophic traits variation with metabarcoding data. This study illustrates how metabarcoding data obtained from feces can be combined with modelling and simulation approaches to test hypotheses in the conventional analytic framework of trophic analysis.

## 1 Introduction

Trophic studies are central to our understanding of ecological interactions, providing insights into the processes that structure ecological communities and regulate energy-flow through trophic networks (Garvey & Whiles, 2016; Nielsen et al., 2018). Functional differences among species are thought to be one of the main sources of ecological variability in food webs, and many diet studies have been performed to investigate inter-specific trophic niche variation (Bolnick et al., 2003). However, there is now a growing body of evidence that intra-specific trait variability (ITV) is crucial for maintaining functional diversity in ecosystems, species and populations (Des Roches et al., 2018; Raffard et al., 2019). Intra-specific trait diversity promotes functional complementarity among individuals through niche partitioning, facilitating more efficient use of ecological opportunities over space and time (Bolnick et al., 2011), which in turn may increase individual fitness and population stability (MacColl, 2011). Intra-specific trait variability can be partitioned into three components: within-individual variation, between-individual variation and population-level variation (Albert et al., 2011). In the context of trophic ecology, these three components correspond to the average individual trophic niche width within a population (WIC; related to α-diversity), the average trophic variation between individuals (BIC; related to β-diversity) and the total trophic niche width (TNW; related to γ-diversity), respectively, TNW being the sum of WIC and BIC (TNW = WIC + BIC) and BIC being related to individual specialization (IS) (Bolnick et al., 2003; Roughgarden, 1974). According to this framework, many *generalist* populations are in fact composed of specialized individuals with relatively narrow diet breadths (Araújo et al., 2011; Bolnick et al., 2002, 2003).

For predators, ecological opportunity, i.e. the availability of ecological resources that may be exploited at a given moment (Stroud & Losos, 2016; Wellborn & Langerhans, 2015), varies across space and time (seasonality). Environments are subject to temporal changes, which in turn interact with prey phenology (Conover, 1992; Marshall & Burgess, 2015; Merritt et al., 2001). In rivers for instance, the abundance of invertebrates is known to be influenced by flow regime dynamics (Monk et al., 2008; J. C. White et al., 2017), temperature (Arscott et al., 2003; Haidekker & Hering, 2008), substratum composition (Downes et al., 2000), and water chemistry (Cross et al., 2006; Gafner & Robinson, 2007). In parallel, aquatic insects have a complex life history with most having a short, aerial adult period and an aquatic juvenile stage. The periodicity of the juvenile stage can vary from a few months to a few years depending on each species’ reproductive strategy (e.g. voltinism) and local environmental conditions (Clifford, 1982; Corbet et al., 2006; Kong et al., 2019). The diverse and overlapping nature of prey phenologies, and their interaction with environmental temporal changes, can lead to significant seasonal variation in community assemblages and in abundances (Erba et al., 2003; García et al., 2007) and consequently on the availability of prey for fishes.

Empirical studies have confirmed that seasonality is a major source of trophic niche variation (e.g. Falke et al., 2020; Varpe & Fiksen, 2010; Shutt et al., 2020) that has implications for individual fitness (Durant et al., 2005; Hipfner, 2008) and affects the demography of populations (Miller-Rushing et al., 2010). In response to low resource availability, trophic niche expansion is predicted by the Optimal Foraging Theory (OFT; see Perry & Pianka 1997). This niche expansion is achieved by the introduction of novel resources into the population’s niche when preferred resources are less abundant. Trophic ITV has been shown to be a determinant component of this process (Bolnick et al., 2011; Sjödin et al., 2018), notably in the case of seasonal change in ecological opportunities (Costa-Pereira et al., 2017). The relationship between trophic niche variation (especially ITV) and ecological opportunity is therefore central for the understanding of biodiversity evolution (Bolnick et al., 2003, 2011; Brodersen et al., 2018), and for conservation and management issues (Agosta, 2002; Johnson et al., 2009; Titulaer et al., 2017).

For studying trophic niche variation, recent methodological advances in diet analysis by metabarcoding now provides a non-destructive and valuable alternative to methods based on the morphological identification of prey or stable isotopes (Alberdi et al., 2019; Sousa et al., 2019). Notably, metabarcoding offers better taxonomic resolution of prey types, and has proven to be a powerful tool for characterizing complex trophic networks (Clare, 2014; Roslin et al., 2016), the niche partitioning between species (Pansu et al., 2019; Soininen et al., 2015), and for guiding the conservation and management of endangered species (Brown et al., 2014; Quéméré et al., 2013). Here we used a metabarcoding approach to characterize intra-specific trophic niche variation in the fish *Zingel asper* (L.) [Actinopterygii: Perciformes: Percidae]. *Zingel asper* is a critically endangered species (Crivelli, 2006) endemic to the Rhône river basin, that lost roughly 85% of its historical range during the 20^th^ century and is now restricted to five disconnected metapopulations in eastern France and Switzerland (Georget, 2019). *Zingel asper* is a benthic species with a low diel displacement range, that has naturally low population density (Cavalli et al., 2003, 2009; Danancher et al., 2007; Labonne et al., 2003) and mainly feeds on macro-invertebrates (Cavalli et al., 2003; Corse et al., 2019).

Dietary data based on the metabarcoding of *Z. asper*’s feces were used to investigate qualitative and quantitative trophic variation in this species. The Rhône river basin covers an area of ~98 500 km^2^ and contains several climatic zones: northern areas receive the highest precipitation, while southern areas have a Mediterranean climate with hot and dry summer periods (Olivier et al., 2009). Therefore, we investigated the trophic variation in five populations of *Z. asper* that live in representative environmental conditions over the remnant species range. We characterized variation in *Z. asper*’s trophic niche for both individual (WIC and BIC) and population (TNW) trophic traits. In addition to seasonal variation, we tested the effect of two well-known determinants of ITV: body size (a functional trait related to ontogeny) and intra-specific competition (e.g. Bolnick et al., 2010; Svanbäck et al., 2008; Vander Zanden et al., 2000; Zhao et al., 2014). Lastly, changes in predator diet may subsequently affect predator body condition (e.g. Skinner et al., 2016), a life-history trait related to the size of energy reserves of individuals (Peig & Green, 2009) that can drive the future fitness of individuals (Kotiaho, 1999; Wells et al., 2016). We therefore aimed to determine whether trophic niche variation could affect the individual future fitness of *Z. asper* (using body condition as a proxy).

## 2 Materials and methods

### 2.1 Fish and feces sampling

Our sampling scheme included five sites which represent four of the five remnant *Z. asper* populations (see Figure 1). Sampling was performed over several campaigns in spring, summer and autumn in 2014 and 2015 (see Table S1), in accordance with permits from the French *Directions Départementales des Territoires* (DDTs) from *Hautes-Alpes*, *Alpes de Haute Provence*, *Ardèche* and *Jura*. Fishes were caught by electrofishing and then laid in a plastic, wire mesh fishpond until biometrical measures and feces collection were performed. Fishes were weighted (precision 0.1g) and their fork-length measured (Lf; precision 1mm). The abdomen of *Z. asper* individuals was then pressed by hand in order to drain out feces. Feces were immediately placed in a 2ml vial containing 96% ethanol and stored at −20°C. After biometric measures and feces collection, fishes were released within the sampling area. A total of 967 *Z. asper* individuals were caught, and 498 feces samples were collected. Additionally, the surface of the prospected area was measured each sampling season in order to estimate the density of *Z. asper* (which was used as a proxy for intraspecific competition).

**Figure 1.**
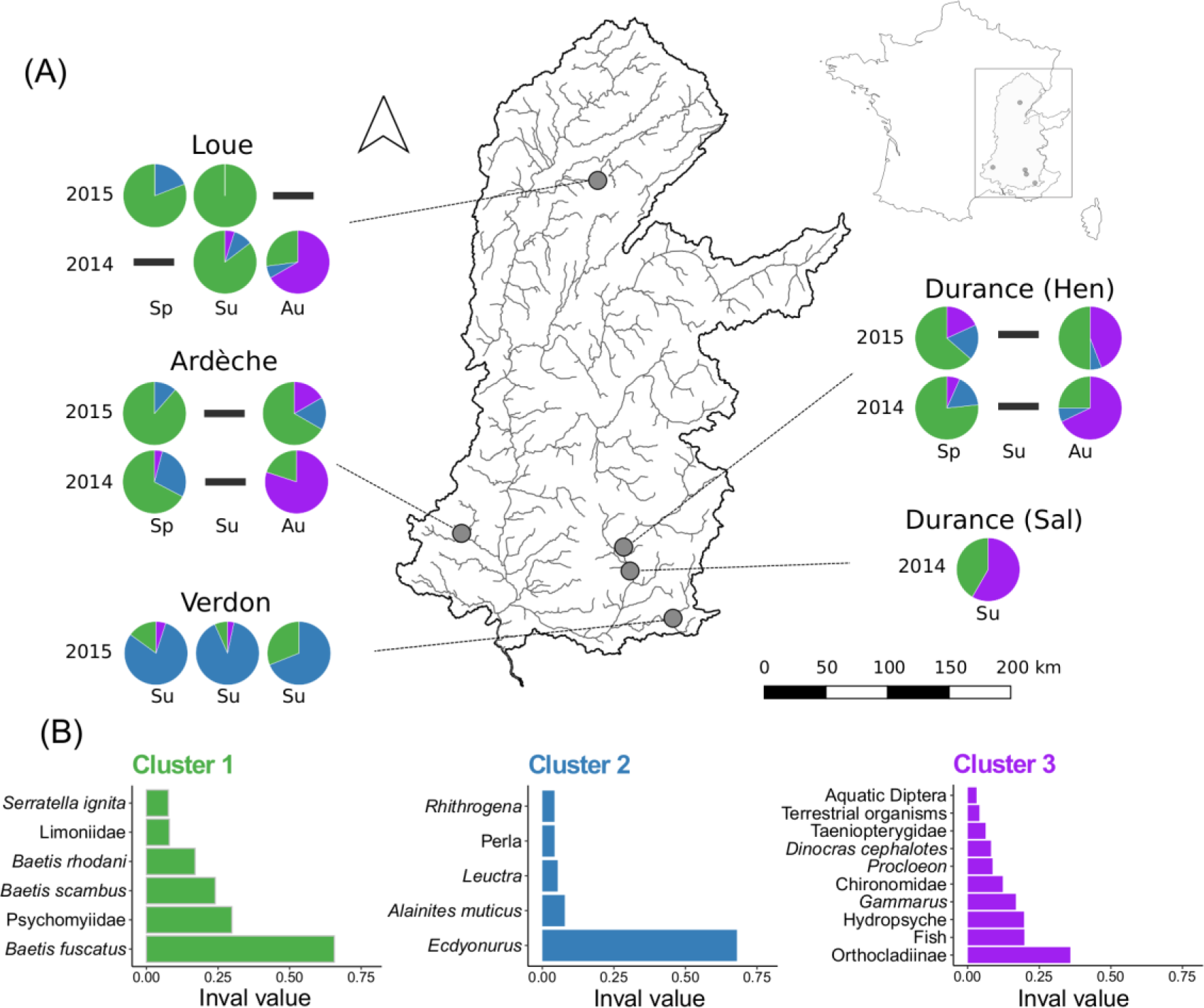
Diets clusters of *Zingel asper* populations (A) and the indicator species value (Indval) histograms for each diet cluster (B).

### 2.2 Diet metabarcoding

Fecal DNA extractions were conducted in a room dedicated to the handling of degraded DNA (*Plateforme ADN Dégradé* of the LabEx CeMEB, Montpellier, France) following the method described by Corse et al., (2017) and the specific safety measures described by (Monti et al., (2015). The DNA was extracted from the whole feces for all individuals. In order to further minimize cross-sample contamination, extraction series were limited to 25 samples, including 21 *Z. asper* feces, two ‘alien’ feces (i.e. from marine or non-European continental predators; see Corse et al., 2019), one negative control for extraction and one negative control for aerosols (for more details on controls, see: Corse et al., 2017, 2019). Additionally, our analyses also included two distinct mock samples communities (described in Corse et al. 2019) as both positive controls and standards across MiSeq runs. Samples and controls were amplified by PCR in triplicates using three primer sets (MFZR, ZFZR and LFCR; Corse et al. 2019) that target ~150bp overlapping sequences located in the 5’ end of the Cytochrome *c* oxidase subunit I gene (COI). Thus, nine separate PCRs were generated per sample/control. Our PCR-enrichment step included the one-locus-several-primers (OLSP) strategy developed by Corse et al., (2019), which aims to minimize false negatives in metabarcoding data by using several primer sets that target overlapping but complementary invertebrate taxa (see also: Esnaola et al., 2018; Hajibabaei et al., 2019). Amplicons were then processed and sequenced on an Illumina MiSeq v3 platform (as detailed in Corse et al. 2017). High-throughput sequencing (HTS) data were filtered using the ASV-centered (Amplicon Sequence Variant) procedure developed by Corse et al. (2017). Several abundance and frequency thresholds were determined from the HTS data of negative and positive controls and from exogenous samples in order to filter out all ASVs that could not be distinguished from low frequency noise (LFN; *sensu* De Barba et al., 2014), thereby minimizing false positives in fecal samples (i.e. experimental/molecular artefacts such as PCR/sequencing errors, tag switching and cross-sample contaminations). We further ensured the reproducibility of ASVs by (i) eliminating PCR replicates considered to be too distant compared to the other replicates from the same sample (Renkonnen dissimilarity used), by (ii) retaining only ASVs that were present in at least two replicates, and by (iii) discarding chimeras and pseudogenes. Ultimately, ASVs obtained from the different primer sets that were identical in their overlapping regions (~130bp) were combined into contigs (further details in: Corse et al., 2017; 2019).

The taxonomic assignment of ASVs/contigs was conducted both automatically using the lowest taxonomic group (LTG) approach (Corse et al., 2017) and manually using BOLD systems (www.boldsystems.org; Ratnasingham & Hebert, 2007). When necessary (i.e. insufficient assignment level, conflicting results between LTG and BOLD), we built phylogenetic trees and/or integrate biogeographic information to finalize taxonomic assignment. In the case of the prey of *Z. asper*, this procedure (which includes three distinct assignment approaches) allowed for species-level identification for up to 75% of the ASVs/contigs (Corse et al., 2017). As the *Z. asper* mainly feeds on macroinvertebrates but can also fed on fishes (Cavalli et al., 2003; Corse et al., 2019; Raveret-Wattel & Bessin, 1900), we considered macroinvertebrates and fishes as relevant prey and collectively referred to them as Macrometazoans. All other taxa (listed in Corse et al. 2019) were excluded from the analyses.

### 2.3 Statistical analyses

All statistical analyses and data formatting were performed in R v 3.5.2 (R Core Development Team, 2018), using the RStudio interface (RStudio, 2010). Prey abundances were estimated using the Minimal Number of Individuals (MNI; White, 1953) approach. MNI is a semi-quantitative statistic that corresponds to the number of distinct ASVs/contigs validated in each sample for a given prey item (see Corse et al. 2017). To evaluate a possible ontogenetic effect on the diet of *Z. asper*, we defined four size-classes: Young of the Year (YOY; Size-class 1), 63-108 mm (Size-class 2), 109-151 mm (Size-class 3), and 152-205 mm (Size-class 4). With the exception of the YOY, the size classes were based on quartiles.

#### 2.3.1 Qualitative trophic variation

*Zingel asper*’s diet was first examined qualitatively, both spatially (sampling sites), and temporally (seasons and year) by conducting (i) a Principal Component Analysis based on proportions (pPCA), performed on the relative abundances (based on MNI) of prey species, and (ii) a hierarchical agglomerative clustering analysis (Ward, 1963). The cluster analysis was performed using the Ward clustering algorithm (function *agnes*, package *cluster*; Maechler et al., 2012) based on dissimilarity measurements calculated from Hill number β-diversities (q = 1, function *pair_dis*, package *hilldiv*; Alberdi and Gilbert 2019a). The optimal number of clusters was determined by the Gap-statistic calculated using the *fviz_nbclust* function from the package *NbCluster* (Charrad et al., 2014). The diet clusters were then characterized by Dufrene-Legendre indicator species analysis (Dufrêne & Legendre, 1997), which calculates an indicator value for each prey item in the respective clusters (function *indval*, package *labdsv*; Roberts & Roberts, 2016).

#### 2.3.2 Estimation of trophic traits

Quantitative dietary niche variation of *Z. asper* was examined for two individual trophic traits: the individual diet niche width (INW; related to α-diversity) and the between individual component of diet niche width (BIC; related to β-diversity). The INW is related to the Within-Individual Component (WIC) of the trophic niche width when using diet data (Bolnick et al., 2002). Both INW and BIC were estimated using ‘traditional’ estimators and using the Hill numbers diversity indices (Hill, 1973). The sensitivity of Hill numbers (^q^D) to rare prey types can be adjusted with the order of diversity parameter q (Alberdi & Gilbert, 2019a). We calculated Hill numbers at q = 1, which includes the relative abundance of prey (in our case, MNI) in the diversity calculation. INW was estimated by the Shannon-Wiener index (INW_S_) (Weaver, 1949) using the R package *vegan* (Oksanen et al., 2008), and by Hill numbers ^1^D (INW_D_) (see Alberdi & Gilbert 2019a), using *hilldiv* (Alberdi & Gilbert 2019b). BIC was first estimated using the individual specialization index V (BIC_V_) (Bolnick et al., 2007) which corresponds to 1 – Psi, where PSi (proportional similarity index) measures the diet overlap between an individual and its population (Bolnick et al., 2002). BIC_V_ was estimated based on the average population diet (pop.diet = “average”) using the *PSicalc* function of the package *RInSp* (Zaccarelli et al., 2013). Secondly, BIC was estimated using dissimilarity measurements derived from β-diversities based on the Hill number ^1^D (BIC_D_) (see above). BIC_D_ is the mean dissimilarity between an individual and the other individuals of its population.

In addition to individual trophic traits, we estimated two population-level trophic traits within the Hill number framework. The total niche with (TNW; related to γ-diversity) was calculated for each sampling campaign as ^1^D using a coverage-based rarefaction approach (Chao & Jost 2012). In order to standardize the TNW estimates between sampling campaigns with different sample sizes, ^1^D rarefaction and extrapolation estimates were conducted at 90% sample-coverage using the function *EstimateD* from the package *iNEXT* (Hsieh et al., 2016). Secondly, prey turnover was calculated for each sampling campaign as an estimate of the overall BIC. Prey turnover was calculated using Jaccard-type turnover (see: Alberdi and Gilbert 2019a) which is calculated from Hill β-diversity values (q = 1; function *beta_dis*, package *hilldiv*). Jaccard-type turnover quantifies the normalized prey turnover rate (across individuals) with respect to the whole system (in our case: sampling campaign) (Chiu et al., 2014).

#### 2.3.3 A proxy for fitness potential: the body condition

We estimated the body condition of *Z. asper* using the scaled mass index 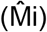, which was calculated using the formula from Peig & Green (2009):

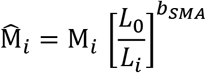

where *Mi* and *Li* are the body mass and length, respectively of individual *i.* L_0_ represents an arbitrary length value; here, we used the mean length value of the fishes caught in this study. *b_SMA_* is the scaling component calculated by standardized major axis (SMA) regression of M on L (Peig & Green, 2009). A one-way ANOVA was performed to confirm that no significant differences in 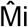 existed between individuals from different size-classes.

#### 2.3.4 The effect of seasonality, intra-specific competition and ontogeny on ITV

To quantify the effect of season, ontogeny and intra-specific competition on INW and BIC, we fitted linear mixed-effects regression models (LMM) using the *lme4* package (Bates et al., 2019). The normality of model residuals was determined visually by q-q plot. The INW_D_ model residuals failed to meet normality, INW_D_ was therefore log-transformed to meet model assumptions. LMMs were fitted using season, size-class and *Z. asper* density as fixed effects and using sampling site and year as random effects. The best performing model was selected based on conditional Akaike information criterion (cAIC), and in cases of very similar cAIC values (differences < 2), the simplest model was selected (function *cAIC*, package *cAIC4*) (Säfken et al., 2018). *P-*values were fitted and calculated for all fixed effects in the best performing LMMs using the package *lmerTest* (Kuznetsova et al., 2017). Model performance was evaluated by conditional R^2^ (R^2^; variation explained including random effects) while fixed effects performance was evaluated by marginal R^2^ (R^2^_M_; variation explained excluding random effects) (function *r.squaredGLMM,* package *MuMIn*; Barton, 2019). The respective contribution of each fixed effect to the marginal R^2^ was determined by type III ANOVA (with Kenward Roger’s degrees of freedom corrections). Differences in traits between the factorial fixed effects were determined by post-hoc Tukey tests on the best performing models using the *emmeans* function with Tukey corrections (function *emmeans,* package *emmeans*; Lenth et al., 2018).

In addition to LMMs, Bayesian regression models (BRMs) were fitted to quantify the effect of season ontogeny and intra-specific competition on INW and BIC. BRMs provide more conservative inferences on biological systems, particularly in cases of pseudo-replication (Lazic et al., 2020). BRMs therefore served to cross-validate the results obtained by LMM. All models were fitted with a Gaussian family distribution with the exception of INWd, which followed a lognormal distribution. BRMs were performed via the package *brms* (function *brm*) with default priors, 3 chains, 20000 iterations and a warm-up of 15000. Model selection was performed by Leave-One-Out cross-validation (LOOic) via the package *loo* (Vehtari et al., 2017). Differences in the response variables between factorial variables (e.g. season) was determined by calculating the probability of selecting a random value from the posterior predictive distribution (P) for a given grouping (e.g. autumn) and it being higher or lower than another value selected from the posterior predictive distribution of another grouping (e.g. spring).

#### 2.3.5 Partionning the variance of TNW

The total niche width of a given population is described as the sum of WIC and BIC (TNW ~ WIC + BIC; Bolnick et al., 2003; Roughgarden, 1972). Determining whether the TNW is mainly driven by variation in BIC and/or WIC provides important insights into the biological mechanisms that drive TNW (Sjödin et al., 2018). We therefore isolated the relative contribution of INW (proxy for WIC) and BIC to variation in TNW using relative importance analysis (package *relaimpo*; Groemping & Matthias, 2018). However, for this type of analysis, the dietary data used to estimate dietary traits should account for a substantial temporal window (MacColl, 2011). If the sampled diet is substantially underestimated due, for instance, to single time-point data and/or limited stomach size, BIC will tend to be overestimated (Bolnick et al., 2007). This can drive artefactual correlations between BIC and TNW. It is therefore necessary to use null models to test whether the observed BIC values are different than what would be expected by random subsampling of the population’s diet (Bolnick et al., 2007). We simulated dietary data using a null model that constrained the number of prey items consumed by each individual to the number in the observed diet data (as described in Bison et al., 2015). Simulations were calculated based on the average diet of each sampling campaign using the package *RinSp*, function *Null.Hp.RInSp* (Zaccarelli et al., 2013). The between-individual component (BICd_simulated_) was then calculated for the simulated diet data using the method described above (section *2.3.2*). In order to test whether an increase in TNW corresponded to a larger increase in BIC than would be predicted under the null model, we compared the slope of (BIC ~ TNW) between observed and simulated BIC values using linear regression. We added an interaction fixed effect (TNW*data type) to test whether the relationship between BIC and TNW significantly differed from the null model. Lastly, we tested for differences between the simulated and observed BIC values for each season using Welch’s t-tests.

#### 2.3.6 The effect of trophic variation and seasonality on fitness potential

To evaluate the effect of dietary variation on fitness potential, LMMs and BRMs were performed following the formula 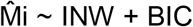, with sampling site and year as random effects. LMMs and BRMs were fitted using ‘traditional’ estimates of INW and BIC, as well as their Hill numbers equivalents. A qualitative estimate of BIC was also included in the model selection process: the dietary cluster assigned to each individual using the hierarchical agglomerative clustering procedure described above. Lastly, to test for seasonal differences in 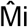, we fitted LMMs and BRMs, again including sampling site and year as random effects. LMMs and BRMs were performed as detailed above (section 2.3.4), including the model selection process (i.e. selection by cAIC and LOOic for LMMs and BRMs, respectively)

## 3 Results

### 3.1 Metabarcoding data

The raw dataset was gathered from 21 distinct MiSeq runs. After pre-processing, the dataset consisted of 61 million (M) reads that corresponded to the three PCR replicates of 510 eDNA samples, 160 negative controls, and 42 mock community samples. Filtering thresholds were determined by run, based on variant occurrence and frequencies (filtering parameters including LFN thresholds are reported in Supporting Information Table S1) and were then applied separately for each run. After filtering, 393 ASVs were validated for MFZR, 447 for ZFZR, and 417 for LFCR. These corresponded to 0.3% of the ASVs initially identified as COI and 70.7% of the reads identified as COI (43 M validated reads). After combining the MFZR, ZFZR, and LFCR ASVs, 391 contigs and 356 ASVs were obtained. At the end of the filtering process, 12 of the initial eDNA samples did not contain any validated ASV or contigs. None of the negative controls had validated ASVs or contigs. All ASVs expected in mock samples were retrieved. As previously reported (Corse et al. 2019), one or two extra ASVs were also validated in most of our mock samples (contig_0238, contig_0124, MFZR_000591; Supporting Information Table S2). By recovering all of the expected taxa from the mock communities in all of the different runs, we assumed that we had minimized random fluctuations, making our samples comparable between runs (see Bakker 2018). A total of 745 distinct Macrometazoan ASVs/contigs (corresponding to 141 prey items) were obtained from the 498 *Z. asper* feces. About 70% of the prey items were identified to the species level, another ~20% to the genus level, ~8% to the family level and ~2% to the order level (Supporting Information Table S2).

### 3.2 Spatial and seasonal qualitative diet variation

The most important prey group for *Z. asper* were Ephemeropteran species (Figure S1): 60% of the total prey abundance (estimated using the MNI), and occurrence in 86% of all fecal samples. Baetidae was the most represented Ephemeropteran family (39% of the total prey abundance), with a single species (*Baetis fuscatus*) representing 28% of the total prey abundance and being present in 70% of fecal samples. The pPCA analysis (Figure S2) identified three main prey types that characterized spatial and temporal variation in the diet of *Z. asper*: PC1 was primarily characterized by the Ephemeroptera genus *Ecdyonurus* (64.0%) while PC2 was mainly characterized by *B. fuscatus* (55.3%) and Orthocladiinae (Diptera) (23.6%). During spring and summer, the diet of *Z. asper* was primarily composed of Ephemeroptera larvae (*B. fuscatus* and *Ecdyonurus*). In autumn, Ephemeroptera consumption declined while the consumption of secondary (e.g. Chironomidae, *Hydropsyche*) and rare prey types (e.g. Plecoptera, fish) markedly increased (Figures S1 & S2). Spatial diet variation was discernible in summer: in the Verdon River the *Z. asper* diet was dominated by *Ecdyonurus*, while at site SAL on the Durance River, the summer diet more closely resembled the autumn diet of the other sampled locations. However, the SAL sampling campaign was almost exclusively composed of feces from Young Of the Year (YOY) individuals (see Table S1).

Hierarchical cluster analysis revealed three distinct diet clusters (Figure 1; Figure S3). The clusters were largely characterized by a single or a few prey-types (see Figure 1): Cluster 1, by *B. fuscatus* (Indval = 0.66), Cluster 2 by *Ecdyonurus* (Indval = 0.68) and Cluster 3 by Orthocladiinae, fish and *Hydropsyche* (Indval = 0.36, 0.20 and 0.20, respectively) as well as several other rare prey. The clustering analysis indicated that the diet of *Z. asper* diversified in autumn when compared to spring and summer, this pattern being consistent among the different sampling sites and between the two sampling years (Figure 1). In spring and summer, Cluster 1 (*B. fuscatus*) was the dominant cluster (except in the Verdon River, where Cluster 2 was dominant). In autumn, the diet of *Z. asper* diversified: prey from Cluster 1 declined, and prey from Cluster 3 (Orthocladiinae and rare prey) proportionately increased. Moreover, in accordance with the pPCA results, the clustering analysis indicated that the YOY differ from the 1yr + individuals in their diet. Most YOY diets were assigned to Cluster 3, irrespective of season (see Figure S3), while Cluster 3 accounted for only ~ 5% to 10% of diets in the other size-classes.

### 3.3 *Individual trophic trait variation in* Zingel asper

Variation in individual traits (Individual Niche Width; INW and Between-Individual dietary Component; BIC) was examined using traditional indices (INW_S_ and BIC_V_) and their Hill numbers equivalents (INW_D_ and BIC_D_). In both cases, the corresponding traditional and Hill number estimates were highly correlated: for BIC_V_ vs. BIC_D_, R^2^ = 0.91 (*p* < 0.001); and INW_S_ vs. log (INW_D_), R^2^ = 1, (*p* = 0; Figure S4).

According to LMMs, roughly 16% (for INW) and 40% (for BIC) of the observed variance was explained by fixed effects alone. Both INW indices were jointly explained by density (intra-specific competition) and size-class fixed effects (Table 2). Density was negatively associated with INW, while INW increased incrementally between size-classes. According to Sum of Square analysis, density and size-class fixed effects explained roughly half of the total variance explained by the best LMM (55% and 45%, respectively) (see Table S3). Post-hoc Tukey tests on model *emmeans* revealed that the differences between size-classes were only significant between YOY (size-class 1) and size-classes 3 and 4 (*p* = 0.006 and 0.007, respectively). In accordance with the LMM results, BRM posterior predictive distribution sampling indicated a 60% and a 64% probability that INW was lower in YOY individuals compared to size-classes 3 and 4, respectively. For the most part, LMM and BRM parameter estimates were in accordance. However, contrary to the best LMM, the best performing BRM also included a seasonal effect. BRM estimates indicated that there was respectively a 60% and 59% probability that INW was lower in autumn compared to spring and summer.

**Table 1.** Glossary.

| Term | Explanation |
| --- | --- |
| Trait | Any measurable feature of an organism e.g. body condition, diet. |
| Intra-specific Trait Variability (ITV) | The individual-based variability of a given trait within a species or population |
| Total trophic Niche Width (TNW) | The total realized niche of a given species or population in terms of prey species richness and abundance. Related to γ-diversity. |
| Individual-Niche-Width (INW) | An individual trophic trait relating to the richness and abundance of prey species consumed by a given individual. Related to α-diversity. Also referred to as the Within-Individual Component (WIC) in other studies. |
| Between-Individual Component (BIC) | An individual trophic trait related to the average difference between an individual's trophic niche and the other trophic niches observed within a given species or population. Relating to inter-individual dietary variability (β-diversity). |
| Prey turnover | A population-level trophic trait indicating the degree of inter-individual dietary variation in a given population or species. |
| Body Condition | Related to the potential fitness of a given individual as measured by the ratio between length and weight. Body condition was measured using the Scaled-mass index ($\hat{M}_i$) in this study. |

**Table 2.**
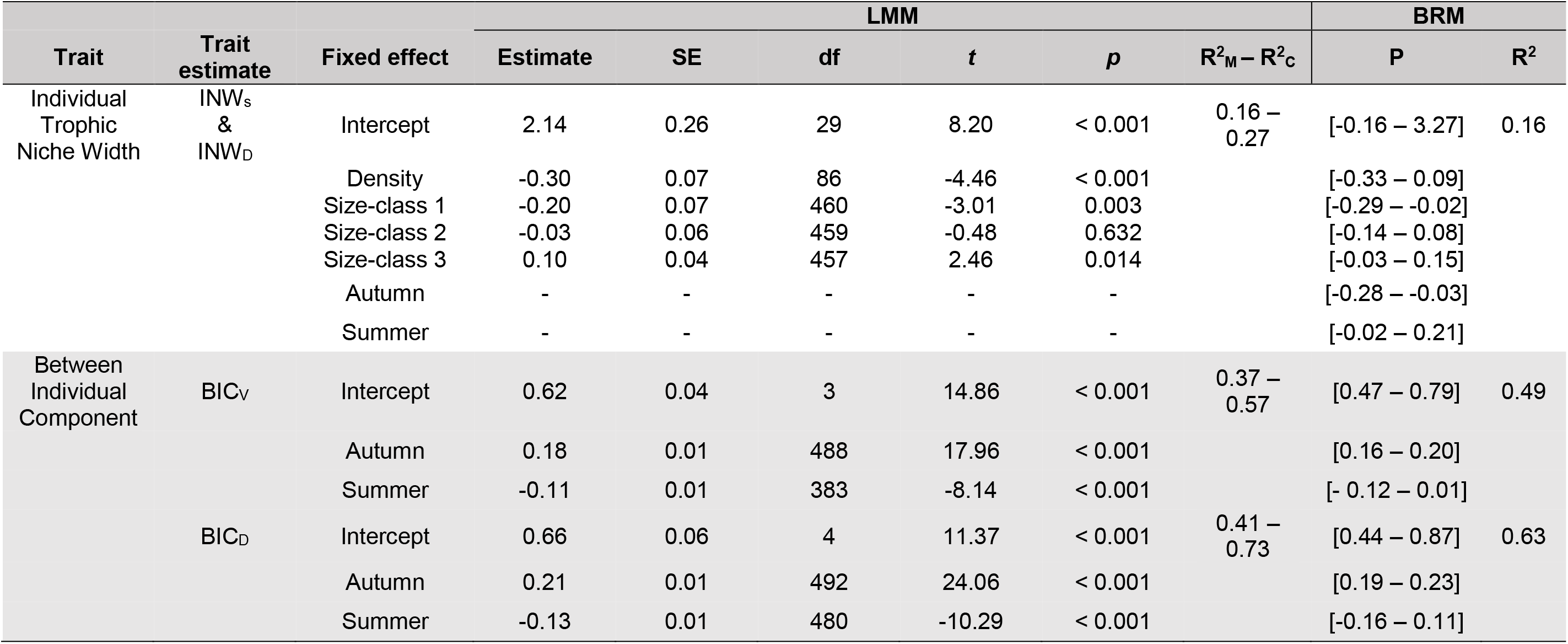
Effect of season, ontogeny and intra-specific competition on *Zingel asper*’s individual dietary traits. SE, standard error; df, degrees of freedom; *t*, *t*-value; *p*, *p*-value; R^2^_M_, marginal R^2^; R^2^_C_, conditional R^2^; P, Bayesian posterior distribution of parameter estimates.

| Trait | Trait estimate | Fixed effect | LMM |  |  |  |  |  | BRM |  |
| --- | --- | --- | --- | --- | --- | --- | --- | --- | --- | --- |
| | | | Estimate | SE | df | *t* | *p* | $R^2_M - R^2_C$ | P | $R^2$ |
| Individual Trophic Niche Width | INW<sub>s</sub> & INW<sub>D</sub> | Intercept | 2.14 | 0.26 | 29 | 8.20 | < 0.001 | 0.16 – 0.27 | [-0.16 – 3.27] | 0.16 |
|  |  | Density | -0.30 | 0.07 | 86 | -4.46 | < 0.001 |  | [-0.33 – 0.09] |  |
|  |  | Size-class 1 | -0.20 | 0.07 | 460 | -3.01 | 0.003 |  | [-0.29 – -0.02] |  |
|  |  | Size-class 2 | -0.03 | 0.06 | 459 | -0.48 | 0.632 |  | [-0.14 – 0.08] |  |
|  |  | Size-class 3 | 0.10 | 0.04 | 457 | 2.46 | 0.014 |  | [-0.03 – 0.15] |  |
|  |  | Autumn | - | - | - | - | - |  | [-0.28 – -0.03] |  |
|  |  | Summer | - | - | - | - | - |  | [-0.02 – 0.21] |  |
| Between Individual Component | BIC<sub>V</sub> | Intercept | 0.62 | 0.04 | 3 | 14.86 | < 0.001 | 0.37 – 0.57 | [0.47 – 0.79] | 0.49 |
|  |  | Autumn | 0.18 | 0.01 | 488 | 17.96 | < 0.001 |  | [0.16 – 0.20] |  |
|  |  | Summer | -0.11 | 0.01 | 383 | -8.14 | < 0.001 |  | [- 0.12 – 0.01] |  |
|  | BIC<sub>D</sub> | Intercept | 0.66 | 0.06 | 4 | 11.37 | < 0.001 | 0.41 – 0.73 | [0.44 – 0.87] | 0.63 |
|  |  | Autumn | 0.21 | 0.01 | 492 | 24.06 | < 0.001 |  | [0.19 – 0.23] |  |
|  |  | Summer | -0.13 | 0.01 | 480 | -10.29 | < 0.001 |  | [-0.16 – 0.11] |  |

**Table 3.**
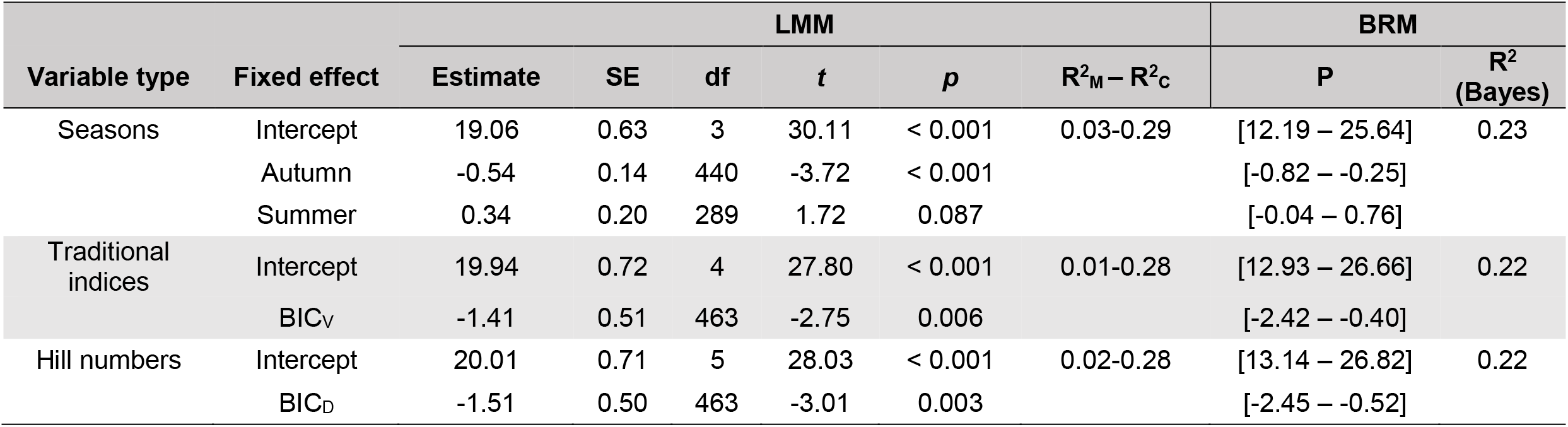
Effect of season and individual diet traits on body condition of *Zingel asper*. Scaled Mass Index 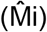 modelled against season and individual dietary traits (INW and BIC). SE, standard error; df, degrees of freedom; *t*, *t*-value; *p*, *p*-value; R^2^_M_, marginal R^2^; R^2^_C_, conditional R^2^; P, Bayesian posterior distribution of parameter estimates.

BIC modelling results revealed that both indices were explained by a seasonal fixed effect, which was retained as the sole predictor for both LMM and BRM models. The seasonal effect was characterized by a significant increase in BIC in autumn compared to both summer and spring (Tukey *p* < 0.001) (Figure 4; Table 2). This autumnal BIC increase was further supported by BRM predictions. For the BICv model, the probability that BIC was higher in autumn than in summer or spring was 90% and 86%, respectively; compared to 93% and 90% for the BIC_D_ model.

**Figure 4.**
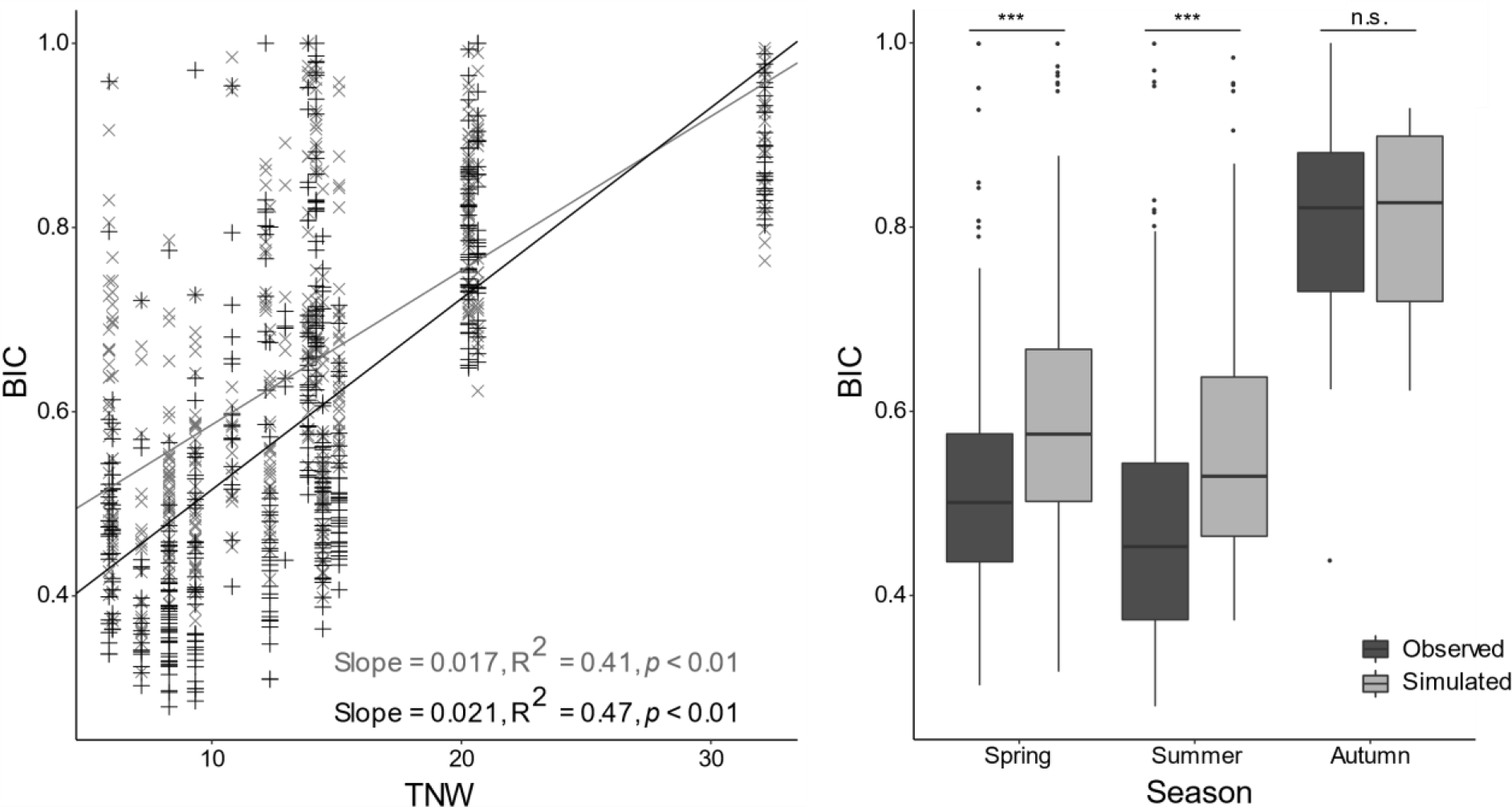
Comparison of observed and simulated BICd values (based on a null model). Left, a scatterplot illustrating the BICd ~ TNW relationship between simulated (grey) and observed BICd values (black). Linear regression slopes are indicated by solid black and grey lines. Right, boxplots comparing simulated BICd values (light grey) to observed BICd values (dark grey) by season. ***, *p* < 0.001; *n.s.,* not significant.

Site and year random effects greatly increased the variance explained for INW indices (R^2^ = 0.16 vs. R^2^ = 0.27). Including sampling site as a random effect also greatly increased the explained variance for both BIC indices (BIC_V_: R^2^_M_ = 0.37 vs. R^2^_C_ = 0.57; BIC_D_: R^2^ = 0.41 vs. R^2^ = 0.73) however, the year random effect failed to improve model fit, indicating negligible annual variation in BIC.

### 3.4 *Populational trophic traits variation in* Z. asper

Total niche width varied from 5.9 to 32.2 across sampling campaigns, with minimum values observed in summer, and maximum values in autumn (see Table S2 and Figure 3). A similar trend was observed in prey turnover, which was generally higher in autumn (from 0.47 to 0.63) compared to spring and summer (from 0.21 to 0.58; see Table S2 for full details). Linear regression models (TNW ~ INW + BIC) revealed that TNW was slightly best explained by Hill number estimates, accounting for 52% of the variation in TNW (R^2^ = 0.52, *p* < 0.001), compared to 46% by traditional estimates (R^2^ = 0.46, *p* < 0.001). In both cases however, between-individual variation accounted for vast majority of TNW variation (BICd = 51%; BICv = 43%). While INW had a significant positive effect on TNW according to linear models, it explained a minimal proportion of the variance of TNW (INWd = 1%; INWs = 3%).

**Figure 3.**
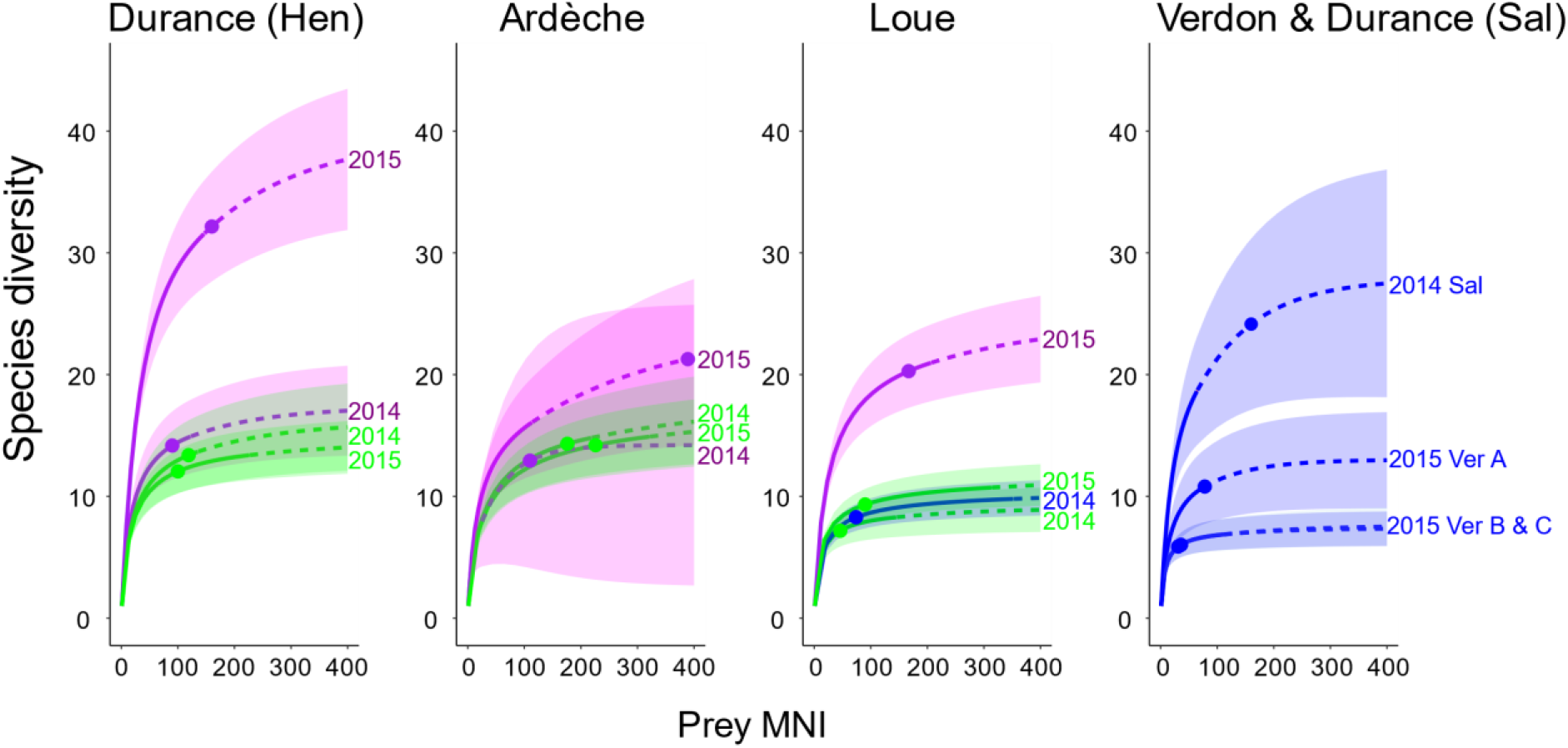
Coverage-based rarefaction curves of Total Niche Width (TNW) by population. TNW corresponds to species diversity calculated using Hill numbers (q = 1), based on 90% diversity coverage. Colors correspond to season (spring = green, summer = blue, autumn= purple).

### 3.5 Comparison between simulated and observed BIC values

Simulated and observed BIC were both strongly correlated with TNW values (Figure 4) however, the slope of the BIC_observed_ ~ TNW regression was significantly greater than for the simulated BIC values (TNW*data type interaction *t* = −2.97, *p* = 0.003). T-tests revealed that simulated BIC values were significantly higher than the observed BIC values in spring (Welch’s t test: *t* = −4.67, *p* < 0.001) and summer (Welch’s t test: *t* = −5.56, *p* < 0.001; Figure 4). However, in autumn observed and simulated BIC did not significantly differ (Welch’s t test: *t* = −0.63, *p* = 0.528).

### 3.6 Determinants of body condition variation

In order to be comparable between individuals of different lengths, body condition 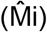 need to account for species-specific asymmetric growth patterns e.g. if shorter individuals are relatively heavier than longer individuals or vice versa. A one-way ANOVA revealed that there were no significant differences in 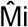 between size-classes in *Z. asper* (*p* > 0.05), indicating that 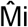 was not biased by the relationship between length and weight in *Z. asper*.

For both Hill numbers and traditional trophic indices, the best performing LMMs and BRMs included a significant negative effect of BIC on 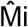 (BIC_D_: *t* = 3.01, *p* = 0.003; BIC_V_: *t* = −2.75, *p* = 0.006) (Table 2). In both cases, neither INW nor dietary cluster was included in the final 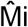 models. According to emmeans performed on the best 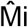 LMM was significantly lower in autumn (spring-autumn: *p* = 0.005; summer-autumn: *p* = 0.018) (Table 2; Figure 2). The BRM also supported a decline in 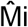 in autumn compared to spring (P = 0.60) and summer (P = 0.62). Overall, individual trophic traits and seasonal variation explained a small proportion of the observed variance in 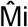 (R^2^_M_ ranged from 0.01 to 0.03), and higher R^2^_C_ values (0.26-0.29) indicated marked spatial and annual variation in 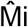.

**Figure 2.**
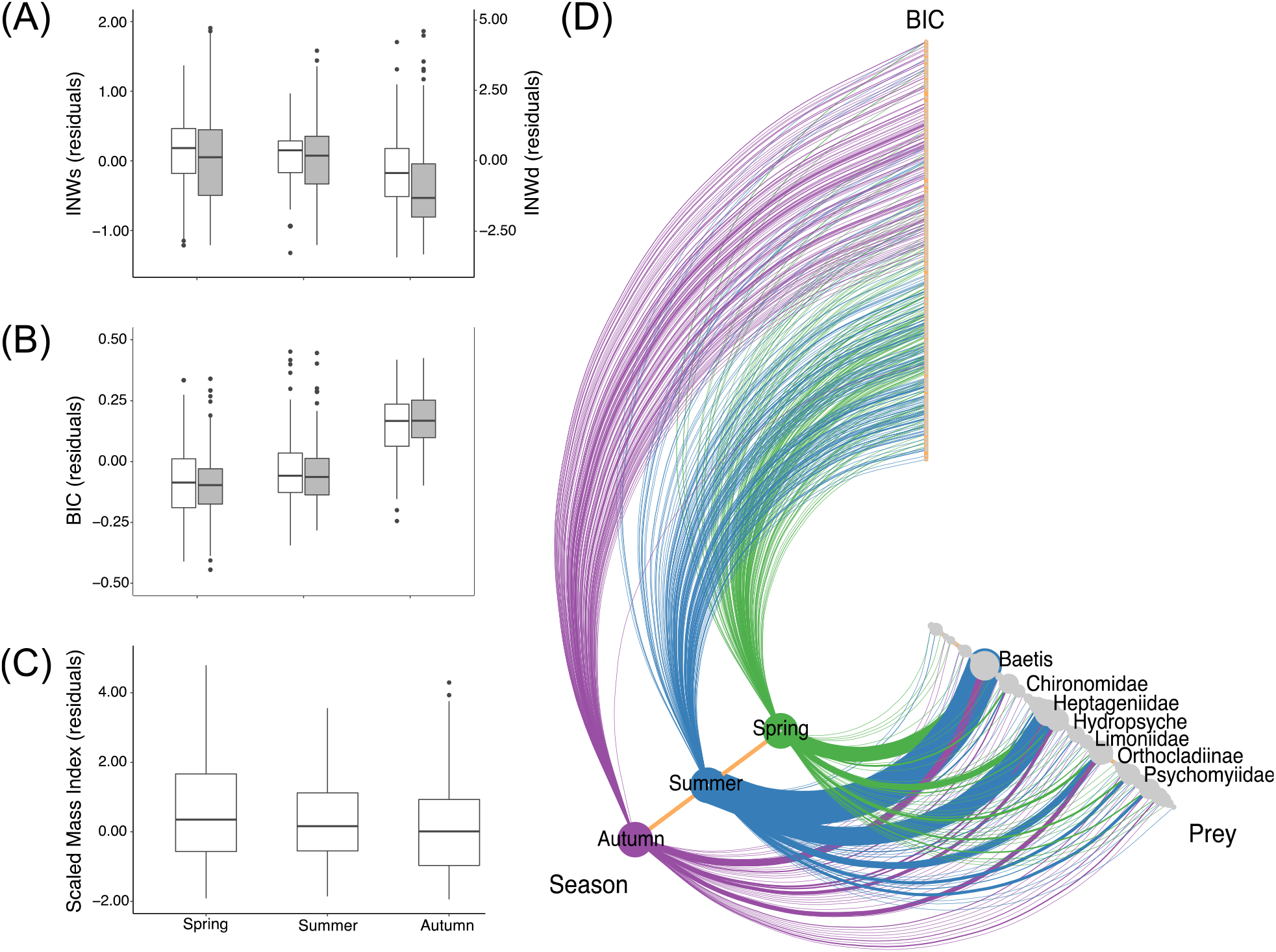
The seasonal variation of INW (A), BIC (B) and 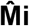 (C), and the network (Hive plot) of relationships of prey, season and *Zingel asper*’s BIC. For INW and BIC boxplots, blank boxes correspond to Shannon and V indices, respectively while shaded plots correspond to hill number indices.

## 4 Discussion

We implemented a robust metabarcoding protocol to characterize individual and population trophic niche variation in *Z. asper*. This protocol used multiple primer pairs to target the COI region to maximize the taxonomic coverage of prey and used an ASV-based filtering procedure that explicitly integrates negative and positive controls to validate dietary metabarcoding data within and among sequencing runs. In accordance with previous local-scale studies (Cavalli et al., 2003; Corse et al., 2019), we found that the *Z. asper* diet was characterized by Ephemeroptera species, closely resembling the diet of its sister species *Romanichthys valsanicola* (Găldean et al., 1997). Additionally, we found qualitative spatial dietary variation of *Z. asper* to be relatively sparse, with *Baetis fuscatus* (Baetidae) and the genus *Ecdyonurus* (Heptageniidae) constituting the two main prey items regardless of the sampling location. The only notable spatial variation was observed in the Verdon River, where the diet of *Z. asper* was characterized by high consumption of *Ecdyonurus* compared to other populations.

### 4.1 Evidence for an ontogenetic trophic niche shift

Cluster analysis revealed a clear distinction between the diet of Young Of the Year (YOY) individuals and 1yr + individuals (size-classes 2-4) (Figure S3). YOY diets more closely resembled the autumn diet of adults (Cluster 3), with higher consumption of Chironomidae. While qualitative dietary variation between size-classes 2 to 4 was minimal, we detected a general positive association between size-class and INW, with the lowest INW observed in YOY individuals. Such dietary niche expansion associated with ontogeny is common in many fish species, owing to size-related limitations in prey consumption, differential nutritional requirements, differences in habitat use or to avoid competition with adult individuals (for a review see: Sánchez-Hernández et al., 2019). Interestingly, adult individuals appear to undergo a seasonal “back-shift” in autumn. Adult individuals both had lower INW and a higher proportion of Cluster 3 diets in autumn (Figure S3), closely resembling the YOY diet.

### 4.2 Seasonal individual trophic trait variation as adaptive foraging?

We found evidence of a seasonal diet shift in autumn that was shared by several *Z. asper* populations in distinct climatic areas of the Rhône river basin. This autumnal shift was associated with a decline in the consumption of *Z. asper*’s main prey type (*B. fuscatus*) and an increase in the consumption of secondary prey (Figure 1). This qualitative dietary diversification coincided with both a TNW expansion and an increase in inter-individual dietary variation (BIC) in most populations (Figures 2 & 3). A seasonal dietary shift was previously reported in the Durance River and was associated with a decline in the abundance of *Z. asper*’s main prey (Baetidae) (Cavalli et al., 2003). Though in our study we did not conduct invertebrate inventories to evaluate prey abundance, it is very likely that the diet shift observed in the Beaume River and Loue River was also driven by the seasonal depletion of *Z. asper*’s main prey species (i.e. *B. fuscatus* and *Ecdyonurus*). Indeed, Trophic Niche Variation (TNV) is driven, among other factors, by variation in ecological opportunity (i.e. the abundance and diversity of prey species) (Varpe & Fiksen, 2010; Shutt et al., 2020; Araújo et al., 2011), and the Optimal Foraging Theory predicts that when the availability of preferred prey species declines, previously disregarded or less preferred prey species will be incorporated into the diet, thus leading to trophic niche expansion at the population level (Perry & Pianka, 1997).

In terms of ITV, we found that the autumnal TNW expansion was associated with a marked increase in inter-individual dietary variation (BIC) (Figure 2b). Transient ITV is a foraging response often related to fluctuations in ecological opportunity (e.g. Brooke McEachern et al., 2006; Matich et al., 2011; Woo et al., 2008), in particular, BIC has been shown to increase as preferred prey declines (Pires et al., 2013; Tinker et al., 2008). BIC is often related to the degree of individual specialization in a given population (Bolnick et al., 2003; Roughgarden, 1974; Thoday, 1974), which can arise due to inter-individual variation in prey preference, microhabitat specialization or individual variation in foraging behavior (Lewis, 1986; Persson, 1985; Svanbäck & Bolnick, 2005). However, as BIC is inherently correlated with TNW it is important to compare the observed BIC against simulated values (based on a null model) to determine whether the observed individual specialization is significant (Bison et al., 2015; Bolnick et al., 2003). While we found that the observed BIC~TNW relationship in *Z. asper* differed significantly from the null (random foraging) model, we found that the observed and simulated BIC values did not significantly differ in autumn (Figure 4). This suggests that the observed autumnal BIC increase arose due to random foraging from a larger spectrum of prey (i.e. high TNW in autumn; Table S4). According to these results, *Z. asper* individuals appear to adapt their foraging behavior towards opportunistic foraging as opposed to specialization when faced with the autumnal depletion of their preferred prey.

Intra-specific competition is also described as one of the primary drivers of trophic Individual Trait Variation (ITV). Intra-specific competition is expected to reduce resource availability thus leading to an increase in INW as individuals expand their diet to include less favorable prey species (MacColl, 2011). However, we found that *Z. asper* density was negatively associated with INW and had no observable effect on BIC. Nevertheless, these unexpected results should be interpreted cautiously due to the co-linearity between season and density (the highest *Z. asper* densities were found in autumn; Table S1) which may have influenced the inclusion of density in the best performing BRM and LMM models. Moreover, model performance (as per R^2^ and AIC) did not greatly differ between models that included both density and season and models that included them as solo predictors, indicating that the variance of INW that is explained by density and season overlap to a degree that it is difficult to disentangle their effects. Given that *Z. asper* is naturally found at low density (e.g. Labonne, Allouche, and Gaudin 2003), we assume that the role of intra-specific competition in driving trophic ITV of *Z. asper* is most likely weak.

Lastly, using the scaled mass index 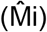 as a proxy for the body condition, we aimed to determine whether the observed autumnal trophic niche expansion constituted a period of trophic stress for *Z. asper*. Body condition incorporates long-term ecological information that ultimately determine an individual’s potential fitness, and/or energy reserves (Kotiaho, 1999). Although we found body condition to be lower in autumn than in summer or spring (possibly indicating that the autumnal dietary shift constitutes a minor trophic stress), we found that BIC and seasonal effects only accounted for a small proportion of the observed variation in body condition (R^2^_M_ = 0.01-0.03). This minor effect of individual dietary traits on the body condition suggests that *Z. asper* optimize their foraging strategy to obtain sufficient nutrients when faced with fluctuations in ecological opportunity so that the seasonal change in individual trophic traits appear to only marginally affect the fitness of *Z. asper*.

Overall, our results associated with the autumnal diet shift in *Z. asper* populations are in line with the Optimal Foraging Theory which predicts that when ‘specialist’ predators are faced with the depletion of their preferred prey, they will adopt a more generalist feeding behavior to meet energy requirements (Holt & Kimbrell, 2007; Singer & Bernays, 2003). Our results suggest that *Z. asper* is specialized on a few ephemeropteran prey species (*B. fuscatus* and *Ecdyonurus*) and adapts its foraging by becoming more opportunistic as preferred prey seasonally decline, to maximize energy gain and maintain fitness.

### 4.3 Estimating individual dietary traits with metabarcoding data

The use of metabarcoding data is now widespread in dietary studies. However, metabarcoding data has rarely been used to estimate individual traits for studying inter- and intra-population level of trophic niche variation (but see Bison et al., 2015; Soininen et al., 2015). Trophic ITV and the conventional analytic framework of trophic analysis were initially developed for diet data obtained by stable isotope analysis or from stomach (or gut, or feces) content analysis. Metabarcoding approaches most closely resemble classical morphological analyses of stomach, gut or feces content as they use the same type of samples to identify prey. Compared to morphological analyses, metabarcoding is known to be poorly quantitative and will provide inaccurate estimations of prey biomass in most cases (Lamb et al., 2019; but see Thomas et al., 2016; Vasselon et al., 2018). However, DNA-based analyses provide much higher taxonomic resolution compared to morphological analyses, which are dependent on the visual identification of prey (Jakubavičiute et al., 2017; Zarzoso-Lacoste et al., 2016). Furthermore, DNA metabarcoding may also detect prey that are morphologically unidentifiable due to the level of degradation of prey species, especially in the case of soft-bodies species (Berry et al., 2015; Egeter et al., 2015; Sutela & Huusko, 2000). The estimation of individual dietary traits is known to be sensitive to the number of prey items considered, which is directly related to taxonomic resolution. Consequently, estimates of dietary traits like INW or BIC are expected to be higher when using metabarcoding data. In trophic studies, this may be particularly meaningful when the taxonomic units have functional differences that could affect the predator’s hierarchization of prey (MacColl, 2011).

The most fundamental difference between stable isotope approaches and stomach, or gut or feces content analyses (being morphological or DNA-based) is related to the temporal window accounted for by the dietary data (Novak & Tinker, 2015; Petta et al., 2020). Compared to stable isotope analyses which are very integrative and can recapitulate the diet of an individual over several weeks or months, the metabarcoding approach employed in this study provided a snapshot of the diet of a given individual, likely accounting for one or a few days of feeding (e.g. Corse et al., 2015). Dietary data that integrate such short temporal windows usually contain a limited fraction of the total prey range of a predator (Aizpurua et al., 2018). Consequently, short-term dietary data tend to underestimate the realized INW as more diverse prey can be incorporated over a longer time-period and may also result in the overestimation of BIC which declines over time as individual diets converge (Novak *et al.* 2015). However, stable isotope analyses also have limitations. For instance, stable isotope analyses are highly dependent on a thorough characterization of isotopic spectrum of prey (which may not be known a priori), and the isotopic diversity of predators does not necessarily relate to taxonomic or functional diversity of prey (Young et al., 2014). Furthermore, physiological variation, growth rate, or stress-related factors can drive significant inter-individual variation in isotopic composition irrespective of the diet consumed (Barnes et al., 2007; Gorokhova, 2018; Karlson et al., 2018) that can hinder trophic studies based on stable isotopes.

Nevertheless, although prudence is needed for their interpretation and comparison, stable isotope-based and morphological- or metabarcoding-based analyses provide valuable, albeit different insights into different dimensions of the trophic niche, including the foraging behavior of organisms (Alberdi et al., 2019; Hette-Tronquart, 2019; Petta et al., 2020). In this study, we used metabarcoding data to describe the mechanisms behind the observed autumnal TNW expansion and BIC increase. The interpretation of our results was assisted by comparing the observed dietary traits to simulated values based on a null model (random foraging model). Simulations revealed that the higher BIC values observed in autumn resulted from a shift toward random foraging as opposed to individual specialization as such increases in BIC are classically interpreted (Bolnick et al., 2003; Tinker et al., 2008). The use of simulations therefore appears to be valuable method to test alternative foraging hypotheses when using metabarcoding data.

### 4.4 Estimating individual dietary traits with Hill numbers

In this study, we used both traditional dietary estimators of INW and BIC (Shannon-Wiener index and V) and their Hill numbers-derived equivalents to characterize the trophic ITV of *Z. asper*. Hill numbers have been proposed as a unifying framework for ecological diversity indices, providing methods for calculating taxonomic, phylogenetic and functional diversity indices (Chao et al., 2014). We found that traditional dietary estimators of INW and BIC were highly correlated with their Hill numbers equivalents (Figure S4) and that the best performing LMMs or BRMs were identical in structure regardless of the type of indices used, suggesting that Hill numbers are highly comparable to traditional diversity indices. The only notable difference was that INW and BIC estimated from Hill numbers were often better explained by ecological variables compared to their traditional counterparts. Our empirical results are therefore in line with the recent calls (Alberdi & Gilbert, 2019a; Ohlmann et al., 2019) for the use of Hill number diversity indices to facilitate comparisons between taxonomic (but also phylogenetic and functional) diversity indices in trophic ecology, especially when using metabarcoding-based data (Alberdi & Gilbert, 2019a).

## 5 Conclusion and perspectives

Diet studies have proven to be critical for guiding the conservation and management of species and habitats (e.g. Nunn et al., 2012; Zarzoso-Lacoste et al., 2019). In particular, an understanding of seasonal trophic niche variation is critical in the current context of climate change (Damien & Tougeron, 2019). Using a high-resolution metabarcoding data and both modelling and simulation approaches to characterize dietary variation in *Z. asper* we were able to relate trophic ITV to several ecological drivers, the seasonal driver being the most important and consistent across the species’ range. Our results provide important insights into how a species can modulate individual and population-level dietary traits to adapt to variation in ecological opportunity (seasonality) and optimize its fitness. However, the high site- and annual-specific variability in ITV and body condition emphasize the need to investigate the causal relationships between the prey community (including its phenological variation), the habitat and both dietary and life-history ITV (e.g. body condition, annual growth) of *Z. asper*. This will improve our understanding of how this critically endangered species adapts to variation in ecological opportunity across space and time, thus providing valuable tools for the management and the conservation of this species.

## Acknowledgments

We warmly thank the staff from the French ‘Office Français pour la Biodiversité’ (OFB), Aix-Marseille Université, INRAE Aix-en-Provence, the ‘Conservatoire d’Espaces Naturels Rhône-Alpes’ (CENRA), and the ‘Parc Naturel Régional du Verdon’ (PNRV) for their help with fieldwork. We especially thank Patrick Gindre, François Huger, Daniel Pedretti and Guillaume Verdier (OFB) for their invaluable technical and logistical support with fieldwork, and we are grateful to Marianne Georget (CENRA), Juliette Dejean (Electricité de France, EDF), Laure Moreau (Syndicat Mixte d’Aménagement du Val Durance; SMAVD), Anne Ferment (PNRV) and Mickaël Cagnant (OFB) for their continuous support during the project. This study is part of the French ‘Plan National d’Action en faveur de l’apron du Rhône’ (2012-2016) coordinated by the ‘Direction Régional pour l’Environnement, l’Aménagement et le Logement d’Auvergne-Rhône-Alpes’ and managed by the CENRA. This study was funded by the SMAVD, the ‘Agence de l’Eau Rhône-Méditerranée-Corse’ (AERMC), the Conseils Régionaux de Provence-Alpes-Côte d’Azur, Bourgogne-Franche-Comté and Auvergne-Rhône-Alpes, the ‘Direction Régionale pour l’Environnement, l’Aménagement et le Logement PACA’ (DREAL PACA) and the PNRV. E.C. was supported by a post-doctoral grant from EDF and OFB, and K.V. was supported by a PhD grant from the ‘Ecole Doctorale des Sciences de l’Environnement’ (ED 251; Aix Marseille Université). Data used in this study were produced by the molecular facilities of LabEx CeMEB (platforms ‘ADN Dégradé’ and ‘GenSeq’, Montpellier), CIRAD (Montferrier-sur-Lez) and SCBM (IMBE, Marseille).

## Author contributions

VD and EC conceived and designed the study. EC, VD, GAS and RC conducted the fieldwork. EC, EM and HV did the molecular work and/or bioinformatics. KV performed statistical analyses, with contributions from EC and VD. KV and VD wrote the original draft, and all authors contributed to further writing and editing.

## Data accessibility

Supplementary data were deposited in Dryad (https://doi.org/10.5061/dryad.2ck7120), including the unfiltered HTS data from MiSeq runs used in this paper.

## Notes

### Competing Interest Statement

The authors have declared no competing interest.

