## Supplementary Tables and Figures for "DNA metabarcoding reveals adaptive seasonal variation of individual trophic traits in a critically endangered fish"

### SUPPORTING INFORMATION

**SUPLEMENTARY TABLES:**

**Table S1.** Sampling locations, sample sizes and fish densities

**Table S2.** Variant and contig, sequence data, taxonomic assignments and diet cluster assignment for fecal DNA samples.

**Table S3.** High-throughput sequencing data filtering parameters.

**Table S4.** Individual-based (mean ± SE) and population-based trophic and life-history traits measured in *Zingel asper* populations.

**Table S5.** Summary of mixed linear regression model performance

**SUPLEMENTARY FIGURES:**

**Figure S1.** The relative abundance of prey in the diet of *Zingel asper* cumulated by sampling campaign.

**Figure S2.** Principal Coordinate Analysis based on proportions (pPCA) of *Zingel asper* prey consumption.

**Figure S3.** Summary of hierarchical clustering results

**Figure S4.** Correlation between traditional and Hill number-derived estimates of INW an

**Table S1. Sampling locations, sample sizes and fish densities.** *N.d.,* not determined

| River | Sampling site ID | Coordinates | Sampling campaign ID | Season | Date | No. Fishes captured | Feces samples (YOY feces) | Fish density (individuals.$ha^{-1}$) |
|---|---|---|---|---|---|---|---|---|
| Durance | Hen | N 44°18’54”, E 5°55’31” | 14HenA | Spring | May-2014 | 68 | 33 (0) | 68.9 |
| | | | 14HenB | Autumn | Oct-2014 | 76 | 34 (0) | 109.7 |
| | | | 15HenA | Spring | May-2015 | 49 | 30 (0) | 49.0 |
| | | | 15HenB | Autumn | Nov-2015 | 52 | 29 (6) | 72.2 |
| Durance | Sal | N 44°7’56”, E 5°59’6” | 14Sal | Summer | Jul-2014 | 40 | 12 (11) | n.d. |
| Ardèche | Plt | N 44°27’18”, E 4°16’39” | 14PltA | Spring | Jun-2014 | 35 | 35 (0) | 18.4 |
| | | | 14PltB | Autumn | Oct-2014 | 66 | 6 (5) | 29.5 |
| | | | 15PltA | Spring | Jun-2015 | 49 | 49 (0) | 26.8 |
| | | | 15PltB | Autumn | Oct-2015 | 51 | 32 (17) | 25.3 |
| Loue | Pln | N 47°0’4”, E 5°49’36” | 14PlnA | Spring | Jun-2014 | 24 | 21 (0) | 8.6 |
| | | | 14PlnB | Summer | Sep-2014 | 49 | 49 (0) | 15.8 |
| | | | 15PlnA | Summer | Jul-2015 | 60 | 41 (0) | 16.5 |
| | | | 15PlnB | Autumn | Sep-2015 | 120 | 48 (2) | 35.3 |
| Verdon | Ver | N 43°44’16”, E 6°20’58” | 15VerA | Summer | Jul-2015 | 82 | 20 (0) | 42.7 |
| | | | 15VerB | Summer | Jul-2015 | 95 | 30 (0) | 54.9 |
| | | | 15VerC | Summer | Sep-2015 | 51 | 29 (0) | 28.1 |

**Table S2. Variant and contig, sequence data, taxonomic assignments and diet cluster assignment for fecal DNA samples. (EXCEL FILE)**

**Table S3. High-throughput sequencing data filtering parameters. (EXCEL FILE)**

**Table S4. Individual-based (mean ± SE) and population-based trophic and life-history traits measured in *Zingel asper* populations.** Total niche width (TNW; Hill q = 1) estimates are based on sample-coverage rarefaction (standardized at 90% coverage), *, extrapolated estimates; n.d., not determined

| | | | Trophic traits | | | | | | | Life-history traits | |
|---|---|---|---|---|---|---|---|---|---|---|---|
| | | | INW | | BIC | | Prey turnover | TNW | | Body condition | |
| River | Site | Campaign | $INW_S$ | $INW_D$ | $BIC_V$ | $BIC_D$ | PT | TNW | 95%CI | $\hat{M}i$ | SE |
| Durance | Hen | 14 Spring | 1.04 | 3.46 | 0.65 | 0.68 | 0.44 | 13.39 | [10.64-16.13] | 19.77 | 0.42 |
| | | 14 Autumn | 0.98 | 2.94 | 0.76 | 0.80 | 0.53 | 14.18 | [11.64-16.72] | 20.08 | 0.34 |
| | | 15 Spring | 1.56 | 5.10 | 0.46 | 0.45 | 0.27 | 12.05 | [10.19-13.92] | 18.82 | 0.28 |
| | | 15 Autumn | 1.14 | 4.42 | 0.82 | 0.88 | 0.63 | 32.16* | [27.76-36.57] | 18.76 | 0.35 |
| Durance | Sal | 14 Summer | 1.44 | 4.44 | 0.69 | 0.76 | 0.58 | 24.15* | [17.90-30.40] | n.d. | n.d. |
| Ardèche | Plt | 14 Spring | 1.31 | 4.31 | 0.54 | 0.52 | 0.35 | 14.32 | [11.30-17.33] | 20.26 | 0.39 |
| | | 14 Autumn | 0.76 | 2.76 | 0.53 | 0.63 | 0.49 | 12.94* | [3.98-21.89] | 17.11 | 1.48 |
| | | 15 Spring | 1.34 | 4.29 | 0.52 | 0.52 | 0.30 | 14.21 | [11.88-16.54] | 20.49 | 0.24 |
| | | 15 Autumn | 0.90 | 2.93 | 0.76 | 0.82 | 0.54 | 21.28* | [13.55-29.01] | 19.13 | 0.41 |
| Loue | Pln | 14 Spring | 1.35 | 4.11 | 0.40 | 0.40 | 0.23 | 7.18 | [5.90-8.46] | 18.74 | 0.34 |
| | | 14 Summer | 1.43 | 4.37 | 0.41 | 0.40 | 0.21 | 8.29 | [7.24-9.34] | 18.80 | 0.24 |
| | | 15 Summer | 1.50 | 4.67 | 0.45 | 0.44 | 0.24 | 9.34 | [8.16-10.51] | 17.50 | 0.20 |
| | | 15 Autumn | 1.18 | 3.57 | 0.75 | 0.79 | 0.47 | 20.29 | [17.49-23.09] | 16.63 | 0.21 |
| Verdon | Ver | 15 Summer A | 0.98 | 2.96 | 0.58 | 0.60 | 0.42 | 10.81* | [7.85-13.77] | 18.32 | 0.30 |
| | | 15 Summer B | 0.92 | 2.78 | 0.45 | 0.48 | 0.25 | 5.88 | [4.82-6.94] | 18.70 | 0.33 |
| | | 15 Summer C | 0.93 | 2.76 | 0.47 | 0.47 | 0.25 | 6.02 | [5.01-7.04] | 19.37 | 0.35 |

**Table S5. Summary of mixed linear regression model performance** for Individual Trophic Niche (INW), Between Individual Component (BIC), and body condition (M̂i). Details include fixed effects Sum of Squares (Sum Sq.), % Variance explained[1], degrees of freedom (df), F value and *p* value.

| Trait | Trait estimate | Best LMM model | Fixed effect | Sum. Sq | Var. Exp. (%) | df | *F* | *p* |
|---|---|---|---|---|---|---|---|---|
| Individual Trophic Niche Width | $INW_S$ + $INW_D$ | Density + Size-class | Density | 4.38 | 55 | 92 | 16.51 | < 0.001 |
| | | | Size-class | 3.51 | 45 | 446 | 4.41 | 0.005 |
| Between Individual Component | $BIC_V$ | Season + density | Season | 5.97 | - | 453 | 165.85 | < 0.001 |
| | $BIC_D$ | Season + density | Season | 8.48 | - | 488 | 303.68 | < 0.001 |
| Body Condition | M̂i | Season | Season | 48.74 | - | 379 | 6.93 | 0.001 |

[1] Variance explained is calculated based on fixed effect's Sum of Squares – it indicates the percentage of the marginal $R^2$ ($R^2_M$) explained by a given fixed effect

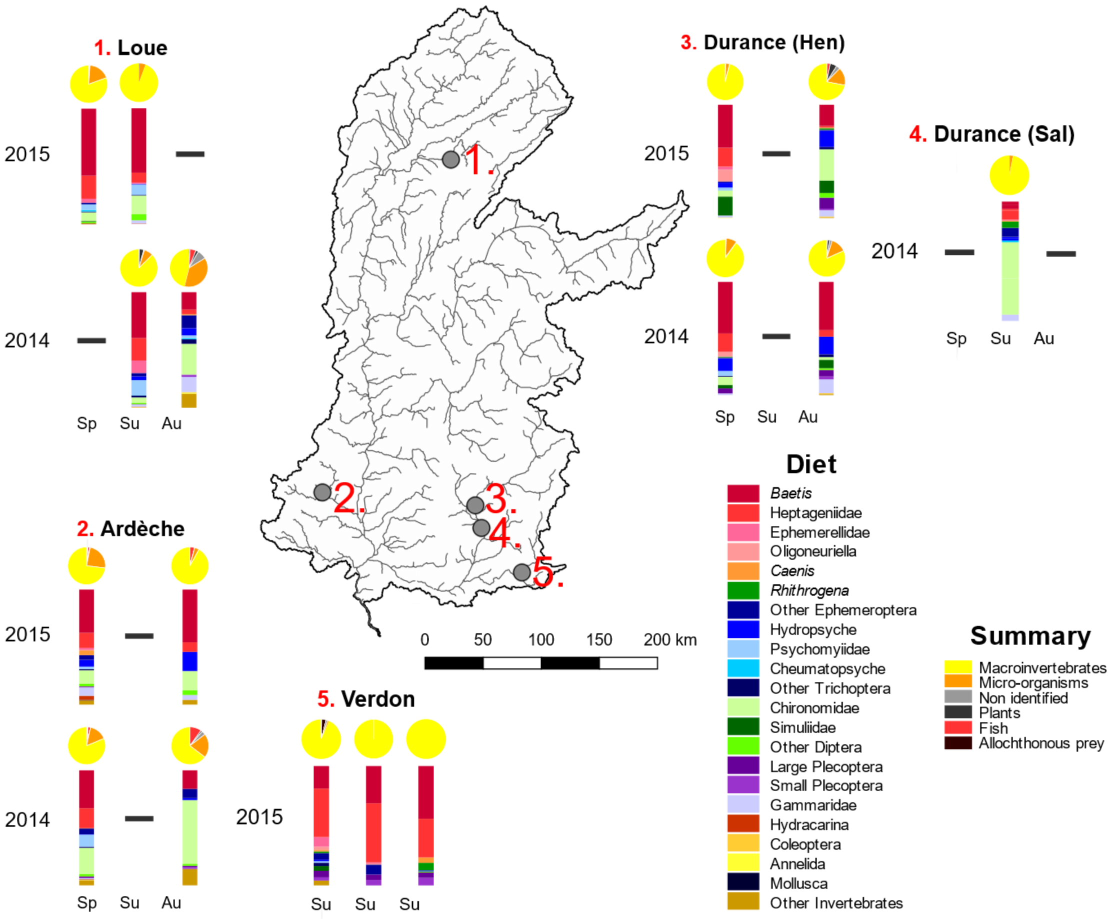


**Figure S1. The relative abundance of prey in the diet of *Zingel asper* cumulated by sampling campaign.** Bar-plots represent the main prey groups of *Z. asper* after the filtration of metabarcoding data. Pie-charts correspond to the proportion of macroinvertebrate, micro-organism, plant, fish, allochthonous and non-identified OTUs prior to filtration. The proportion of prey items are based on the cumulative Minimal Number of Individuals (MNIs).

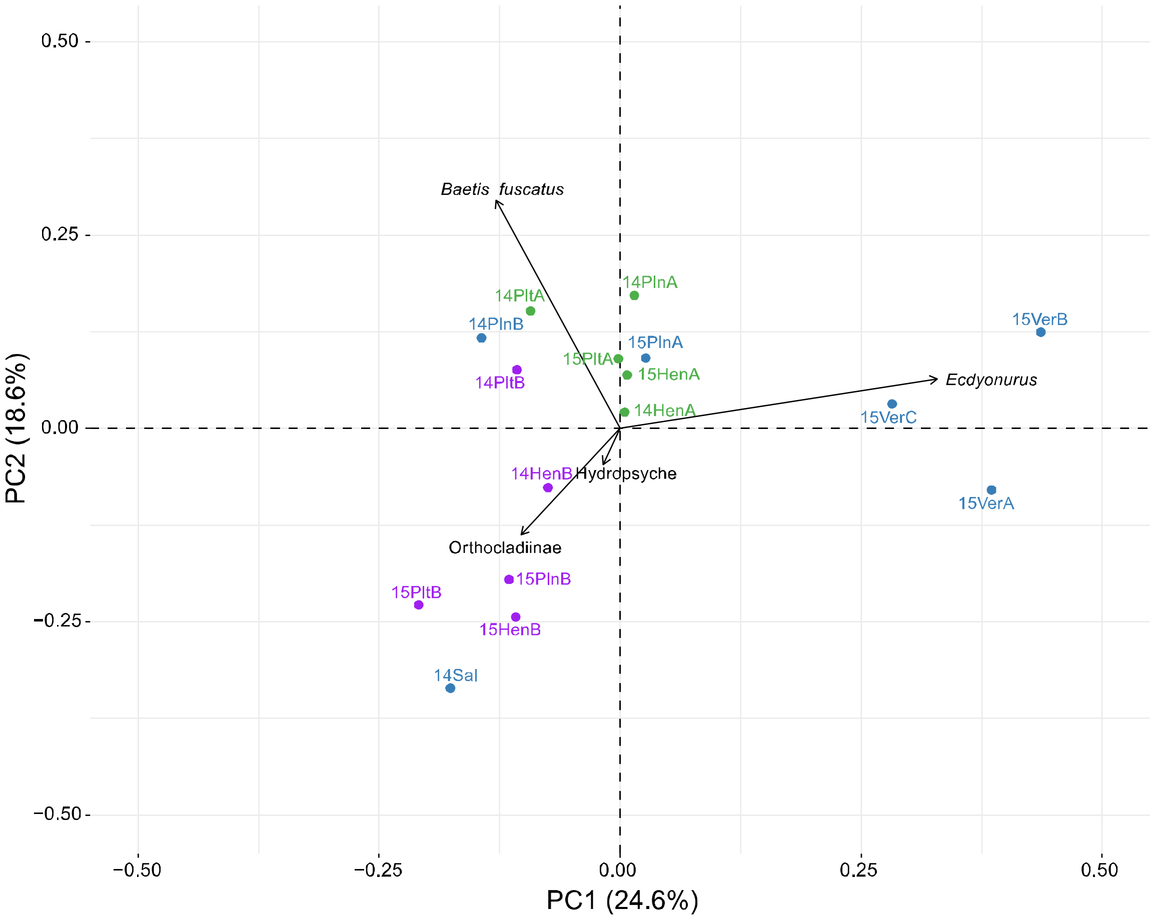


**Figure S2. Principal Coordinate Analysis based on proportions (pPCA) of *Zingel asper* prey consumption.** The proportions of prey items are based on the cumulative Minimal Number of Individuals (MNIs). Each point represents the centroid position of each sampling campaign. Colors correspond to seasons: green = spring, blue = summer and purple = autumn

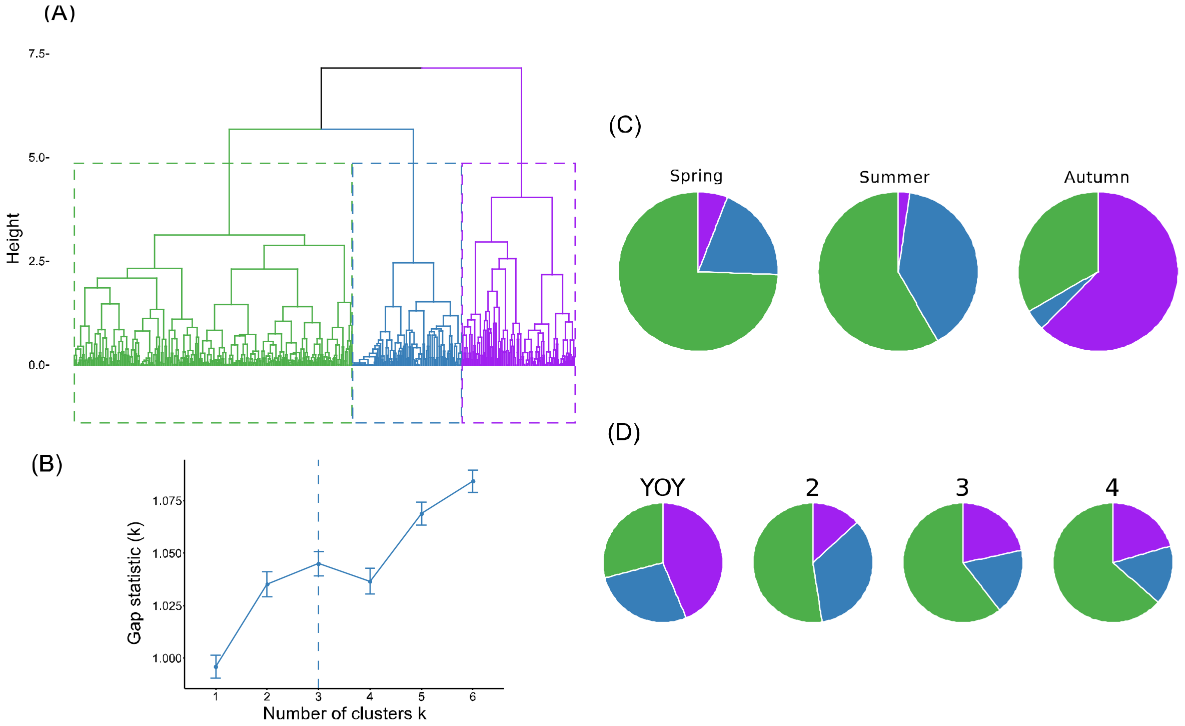


**Figure S3. Summary of hierarchical clustering results**. A) Dendogram, B) Gap statistic results to determine the number of clusters (k) C) Cluster proportions by season (excluding YOY) D) Cluster proportions by size class. Cluster colors correspond to: Cluster 1 (*Baetis fuscatus*) = Green, Cluster 2 (*Ecdyonurus*) = blue and Cluster 3 (Orthodcladiinae and rare prey) = purple.

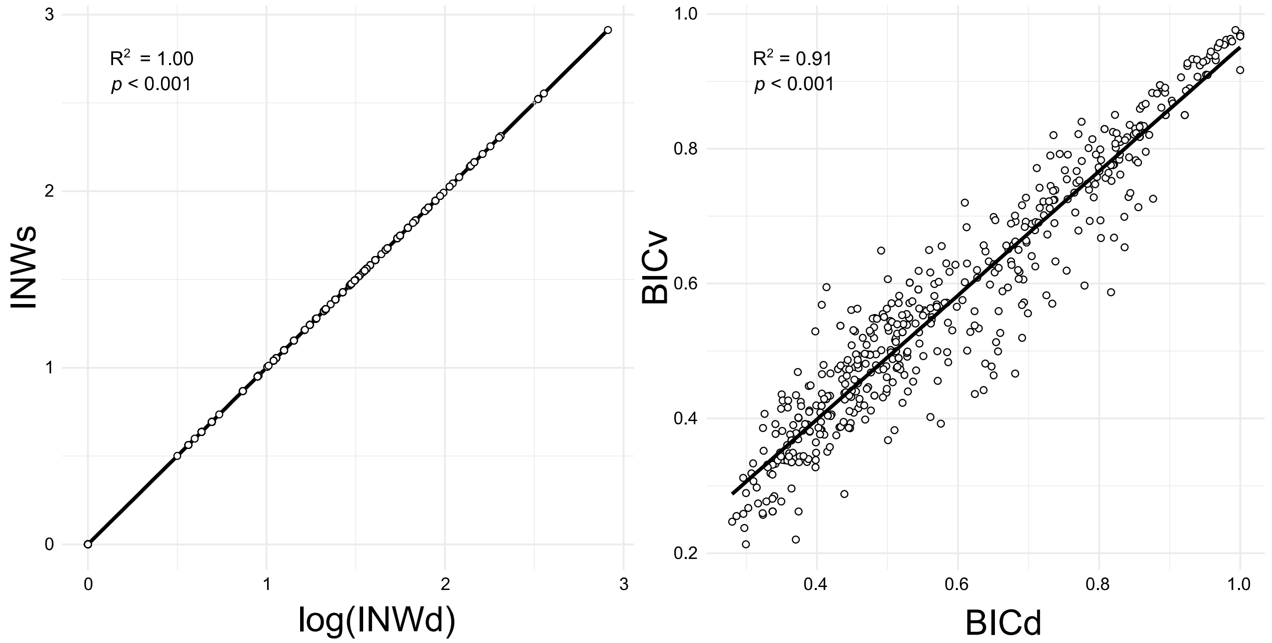


**Figure S4. Correlation between traditional and Hill number-derived estimates of INW and BIC.**
